# Host genetic variation in mucosal immunity pathways influences the upper airway microbiome

**DOI:** 10.1101/062232

**Authors:** Catherine Igartua, Emily R Davenport, Yoav Gilad, Dan Nicolae, Jayant Pinto, Carole Ober

## Abstract

The degree to which host genetic variation can modulate microbial communities in humans remains an open question. Here we performed a genetic mapping study of the microbiome in two accessible upper airway sites, the nasopharynx and the nasal vestibule, during two seasons in 144 adult members of a founder population of European decent. We estimated the relative abundances (RAs) of genus level bacteria from 16S rRNA gene sequences and examined associations with 148,653 genetic variants (linkage disequilibrium [LD] r^2^ < 0.5) selected from among all common variants discovered in genome sequences in this population. We identified 37 microbiome quantitative trait loci (mbQTLs) that showed evidence of association with the RAs of 22 genera (q < 0.05), and were enriched for genes in mucosal immunity pathways. The most significant association was between the RA of *Dermacoccus* (phylum Actinobacteria) and a variant 8kb upstream of *TINCR* (rs117042385; p = 1.61⨯10^−8^; q = 0.002), a long non-coding RNA that binds to peptidoglycan recognition protein 3 (*PGLYRP3*) mRNA, a gene encoding a known antimicrobial protein. A second association was between a missense variant in *PGLYRP4* (rs3006458) and the RA of an unclassified genus of family Micrococcaceae (phylum Actinobacteria) (p = 5.10⨯10^−7^; q = 0.032). Our findings provide evidence of host genetic influences on upper airway microbial composition in humans, and implicate mucosal immunity genes in this relationship.

## INTRODUCTION

Diverse populations of microorganisms inhabit nearly every surface of the human body and these complex assemblies of microbes reflect host-microbe and microbe-microbe interactions. Collectively, these microorganisms constitute the human microbiome (Human Microbiome Project Consortium 2012). Under healthy conditions, the relationship between microbes and the host is symbiotic with many physiologic benefits to the host (Nicholson et al. 2012). Imbalances or changes in the composition of bacterial communities can shift this relationship from symbiotic to pathogenic, a condition known as dysbiosis, which has been implicated in a variety of diseases (Chow et al. 2010). For example, altered composition of airway microbiota has been linked to important respiratory diseases such as sinusitis (Boase et al. 2013), chronic obstructive pulmonary disease (COPD) (Pragman et al. 2012) and asthma (Hilty et al. 2010; Huang et al. 2011; Denner et al. 2015). Similar to the traits it influences, the microbiome itself can be considered a complex phenotype with environmental and genetic factors contributing to its composition (Marsland and Gollwitzer 2014). Understanding how host genetic variation shapes the microbiome, and how the microbiome ultimately functions to modulate host immunity are fundamental questions that are central to fully characterizing the architecture of many common diseases that occur at mucosal surfaces, including those involving the airway.

Although knowledge of the airway microbiome lags behind that of the gut, important characteristics of the microbial communities in the airway are beginning to emerge. Similar to the gut, the community structure of an individual’s airway microbiome is established early in life and plays a critical role in immune development (Arrieta et al. 2014; Gensollen et al. 2016). Many external factors influence the airway microbiome, including mode of delivery at birth (Dominguez-Bello et al. 2010), breastfeeding (Biesbroek et al. 2014), antibiotic use (Noverr et al. 2004; Suárez-Arrabal et al. 2015), and exposure to tobacco smoke (Morris et al. 2013) and pathogens (Bosch et al. 2013). While the influences of environmental exposures on microbiome composition are well known, the degree to which host genetics plays a role in structuring microbial communities is less well understood. In fact, recent data suggest that host genetics may play an important role in shaping microbiome composition. For example, the heritability of the gut microbiome was recently investigated in 1,126 twin pairs (Goodrich et al. 2016). Out of 945 taxa examined, the RAs of 8.8% of taxa had non-zero heritability estimates suggesting that the abundances of those bacteria are influenced by host genetic variation. Moreover, more similar microbiome structures among related individuals compared to unrelated individuals (Yatsunenko et al. 2012; Tims et al. 2013) further supports a role for genetics influencing interindividual variability in microbiome profiles. In fact, quantitative trait locus (QTL) approaches have successfully identified variation in candidate host genes that influence the RA of specific bacteria not only in *Drosophila* and mice but also in humans (Benson et al. 2010; McKnite et al. 2012; Srinivas et al. 2013; Knights et al. 2014; Org et al. 2015; Blekhman et al. 2015; Davenport et al. 2015; Goodrich et al. 2016).

Studies of host genetic influences on the microbiome are particularly challenging due to the profound effects of environmental exposures on microbiome variability. It is not surprising, therefore, that two studies were unable to show host genotype effects on the human gut microbiome (Turnbaugh et al. 2009; Yatsunenko et al. 2012). Studies of related individuals and even twin pairs are confounded to a large extent by the more similar environments among close relatives, making it impossible to completely disentangle the relative roles of genes and environment. To address these challenges, we focused our studies on the Hutterites, a founder population that practices a communal, farming lifestyle that minimizes environmental variation between individuals (Ober et al. 2001), and should increase power to identify genetic influences on complex traits, including the airway microbiome composition. For example, Hutterites prepare and eat all meals in communal kitchens, smoking is prohibited and rare, and individual family homes are nearly identical within each colony (communal farm) and very similar across colonies. Furthermore, the Hutterites in our studies are related to each other in a 13-generation pedigree and are descendants of only 64 founders. Finally, nearly all genetic variation in these individuals has been revealed through whole genome sequencing studies in 98 Hutterite individuals (Livne et al. 2015).

We previously reported studies of the gut microbiome in the Hutterites (Davenport et al. 2014; 2015). Here we interrogated the interaction between host genetic variation and microbiome composition in two accessible sites in the upper airways, the nasal vestibule and the nasopharynx, which have important physiologic functions and relevance to airway diseases. While the nasal vestibule is located in the anterior nares and in direct contact with the environment, the nasopharynx is in the posterior nasal passage and continuous with the lower airway. Overall, our findings demonstrate that the airway microbiome is influenced by host genotype at many loci, and suggest that host expression of innate and mucosal immune pathway genes plays a significant role in structuring the airway microbiome.

## RESULTS

### Nasal microbiome composition

To characterize the variation of the microbiome from the nasal vestibule and the nasopharynx, we first analyzed 16S rRNA V4 gene sequences from 322 samples collected from 144 Hutterite adults in summer and/or in winter months (Table 1). After applying quality control filters and subsampling to 250,000 reads per sample, 83 million reads were assigned to 563 operational taxonomic units (OTUs) with 97% sequence identity. We identified sequences from eleven phyla, with three accounting for 98.94% of the sequences - Firmicutes (52.28%), Actinobacteria (29.81%) and Proteobacteria (16.85%). We then classified OTUs into 166 genera; six dominant genera accounted for 83.30% of the sequences (Figure 1 and Supplemental Table S1).

**Table 1:** Sample composition: A total of 332 samples were collected from 144 (58 male, 86 female) Hutterite adults (age 16 to 78 years).

| Nasal site | Summer | Winter | Both seasons | Unique subjects |
| --- | --- | --- | --- | --- |
| Vestibule | 87 | 80 | 34 | 133 |
| Nasopharynx | 88 | 77 | 40 | 125 |
| Both sites | 72 | 60 | 23 | 144 |

**Figure 1:**
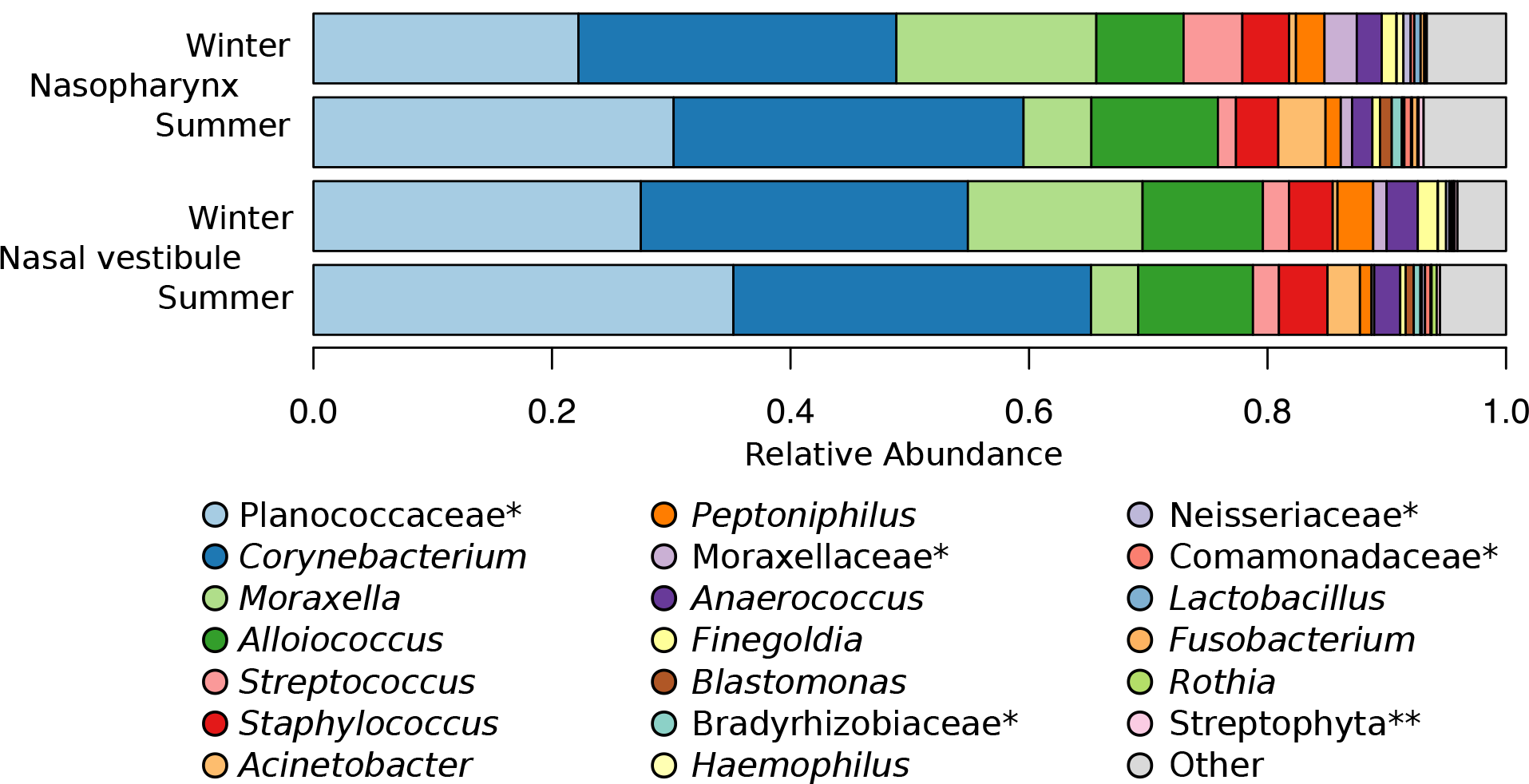
Taxonomic composition of bacterial communities in the nasal vestibule and the nasopharynx, sampled in summer and in winter. Genus level mean RA is shown for the 20 most abundant genera identified in the samples. The remaining 146 genera are grouped as “other”. *Genus unclassified, family level presented. **Genus and family unclassified, order level presented.

In a prior study in a largely overlapping sample of adult Hutterites, we identified large seasonal variation in the gut (fecal) microbiome (Davenport et al. 2014). To see if similar patterns were present in the nasal microbiome, we examined the genus level RAs for individuals studied in both seasons (n=34 for the nasal vestibule and 40 for the nasopharynx). The RA of 12 genera in the nasal vestibule and 15 in the nasopharynx differed by season after applying a Bonferroni correction (paired Wilcox rank sum test, p < 0.0003), nine of which were different between seasons at both nasal sites (Supplemental Table S2). Similarly, we looked for genus level RAs that differed between the nasal sites within each of the two seasons (n=72 for the summer and 60 for the winter) but did not identify statistically significant differences.

### Nasal microbiome diversity

We used three diversity metrics to assess within sample (alpha) diversity for each of the four seasonal nasal site groups - the number of species (richness), Shannon index, and evenness. Overall, the highest alpha diversity was observed in the nasopharynx in the summer (Figure 2 and Supplemental Figure S1A), where the number of observed species and the Shannon index reflected higher diversity compared to the nasopharynx in the winter (paired Wilcoxon signed-rank test; p = 0.002 and 0.048, respectively). Additionally, there was higher diversity in the nasopharynx in the summer compared to the nasal vestibule in the summer (paired Wilcoxon signed-rank test; richness p = 4.6x10^-7^, Shannon index p = 0.009 and evenness p = 0.031, respectively). Although higher diversity trends were observed in the nasopharynx in the summer compared to the winter, these associations were largely due to decreased alpha diversity among women compared to men in the winter (Wilcoxon signed-rank test; richness p = 0.03, Shannon index p = 0.002 and evenness p = 0.001, respectively; Supplemental Figure S1B). Lastly, Shannon index and evenness decreased with increasing age only in the nasopharynx in the summer (p = 0.019; Supplemental Figure S2).

**Figure 2:**
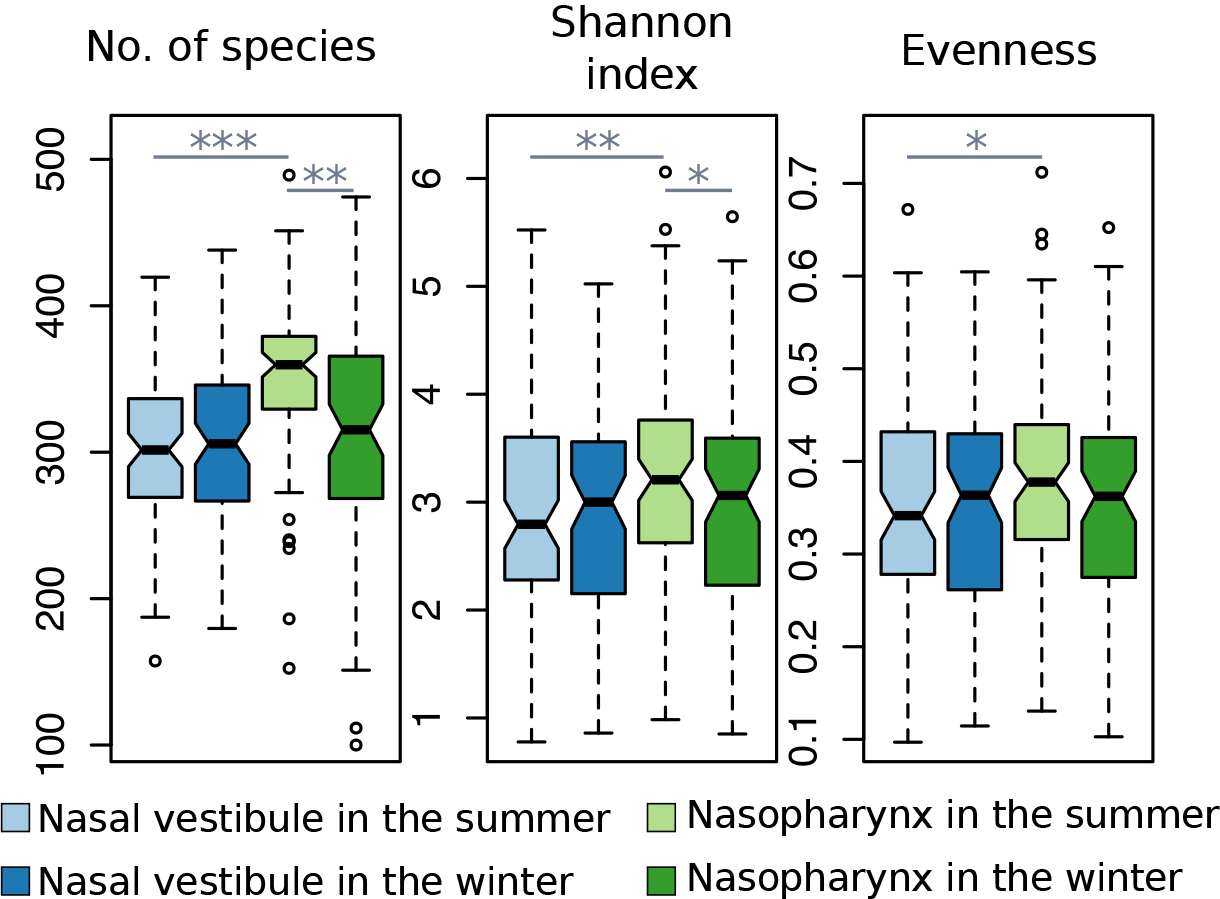
Within sample alpha diversity: Alpha diversity measurements for microbial communities from the nasal vestibule and the nasopharynx by season. The nasopharynx in the summer (light green) shows the overall largest alpha diversity. *Paired Wilcox-rank sum test p < 0.5; ** p < 0.01; ***p < 0.001.

We next analyzed community composition and structure between samples (beta diversity) by calculating Euclidean distances between all pairs of individuals. In the seasonal analyses, the summer samples for both the nasal vestibule and the nasopharynx had lower Euclidean distances compared to their respective winter samples (Wilcoxon signed-rank test, nasal vestibule p < 2.2⨯10^−16^ and nasopharynx p < 2.2⨯10^−16^), reflecting more similar microbiome diversity between pairs of individuals in the summer than in the winter. Moreover, Euclidean distances for the same individual paired with him/herself between seasons (separately within the nasal vestibule and nasopharynx samples) and between nasal sites (separately within the summer and the winter samples) were lower than the respective distances calculated between each individual with all other individuals (Wilcoxon signed-rank test nasal vestibule between seasons p = 9.25⨯10^−8^; nasopharynx between seasons p = 3.97x10^−12^; summer between nasal sites p < 2.2⨯10^−16^, winter between nasal sites p < 2.2⨯10^−16^; Supplemental Figure S3). These results reflect stability in microbiome structure between seasons and nasal sites within individuals, potentially reflecting a genetic component to microbiome composition and diversity.

### Correlation between host genetic similarity and microbiome structure

To evaluate the relationship between genetic similarity (or relatedness) among pairs of individuals and the similarity of their nasal microbiomes, we compared genetic distance, measured by the kinship coefficient and beta diversity between all pairs of individuals in the sample combined across seasons (see methods). We reasoned that if there was a genetic influence on bacterial composition and diversity, more related individuals should have lower measures of beta diversity, reflecting more similar microbiomes. To assess significance we performed 10,000 permutations for each of the two nasal sites. This analysis revealed a significant negative Spearman correlation between kinship and beta diversity (Figure 3).

**Figure 3:**
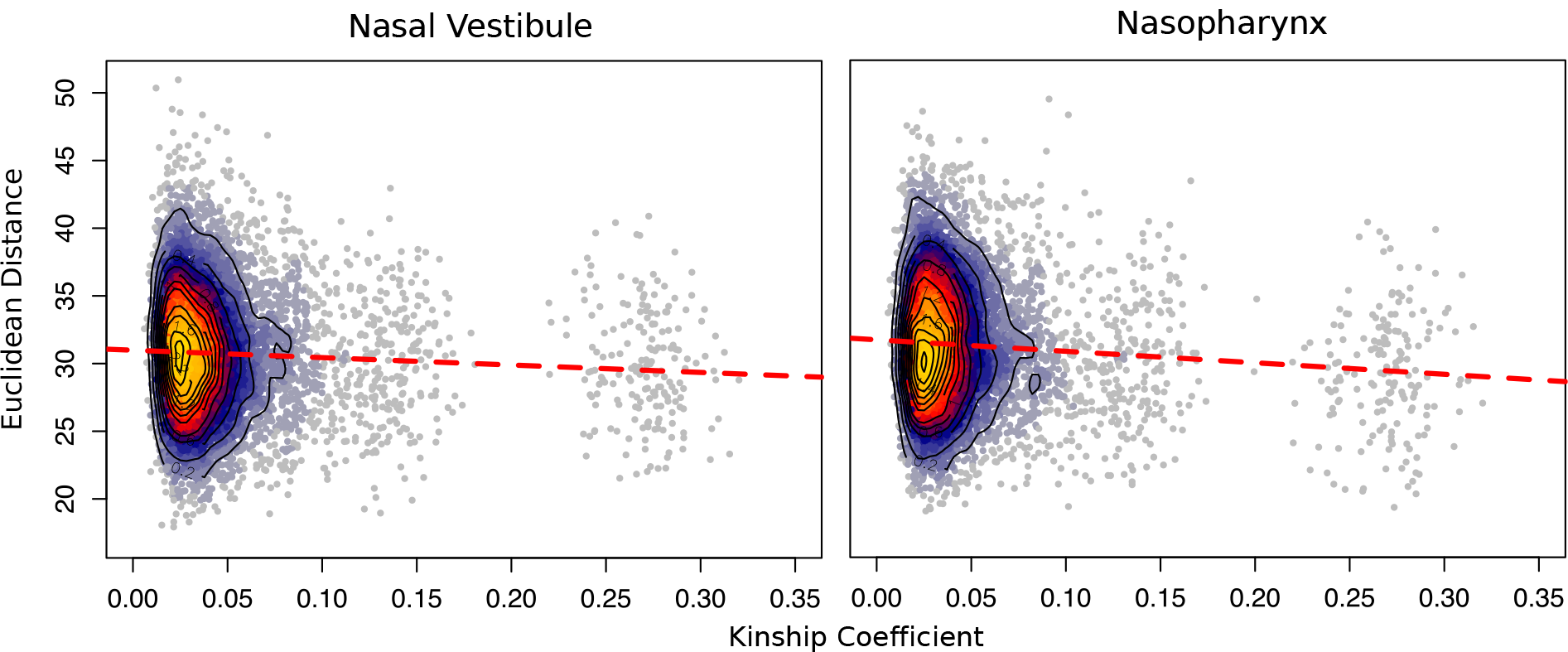
Heat scatterplots of Euclidean distance (beta diversity) by kinship coefficient. Individuals with larger kinship coefficients (more related) have more similar beta diversities (lower Euclidean distances). Red dashed represents the trend line from a linear model. Nasal vestibule p < 1⨯10^−4^; nasopharynx p = 4.0⨯10^−4^.

Although an individual’s microbiome composition is highly sensitive to the household environment (Lax et al. 2014), sharing of households by first degree relatives did not significantly affect the correlation between beta diversity and kinship in our sample. To examine this directly, we removed all first degree relatives who lived in the same household (three sibling pairs and their parents; 15 out of 175 first-degree relative pairs in the sample) and repeated the analysis. The correlation between kinship and Euclidean distance remained significant (nasal vestibule p < 1⨯10^−4^ and nasopharynx p = 5.0⨯10^−4^), indicating that the significant effect of kinship on microbiome similarities between Hutterite adults is not likely due to shared environments. Instead, we attribute these correlations largely to shared genetic variation.

### Genome-wide association studies of relative abundance

To directly test for host genetic effects on genus level bacteria in the nasal vestibule and in the nasopharynx, we performed microbiome quantitative trait locus (mbQTL) mapping on the bacterial RAs for each nasal site in the summer sample and winter samples separately, and in a larger sample combining both seasons. We tested for associations between 52 and 90 genera with 148,653 SNPs (LD r^2^ > 0.5) using a linear mixed model as implemented in GEMMA (Zhou and Stephens 2012), and included sex and age as fixed effects and kinship as a random effect to adjust for the relatedness between all pairs of individuals in our study. Our analyses revealed 37 mbQTLs at q < 0.05, three of which were associated with multiple bacteria (overall 37 variants associated with 22 genus level bacteria). Of the 37 mbQTLs, 14 were associated with 10 genera in the nasal vestibule and 23 were associated with 14 genera in the nasopharynx. The results for mbQTLs with q < 0.05 are shown in Table 2 and results for 108 mbQTLs with q < 0.10 are shown in Supplemental Table S2.

**Table 2:**
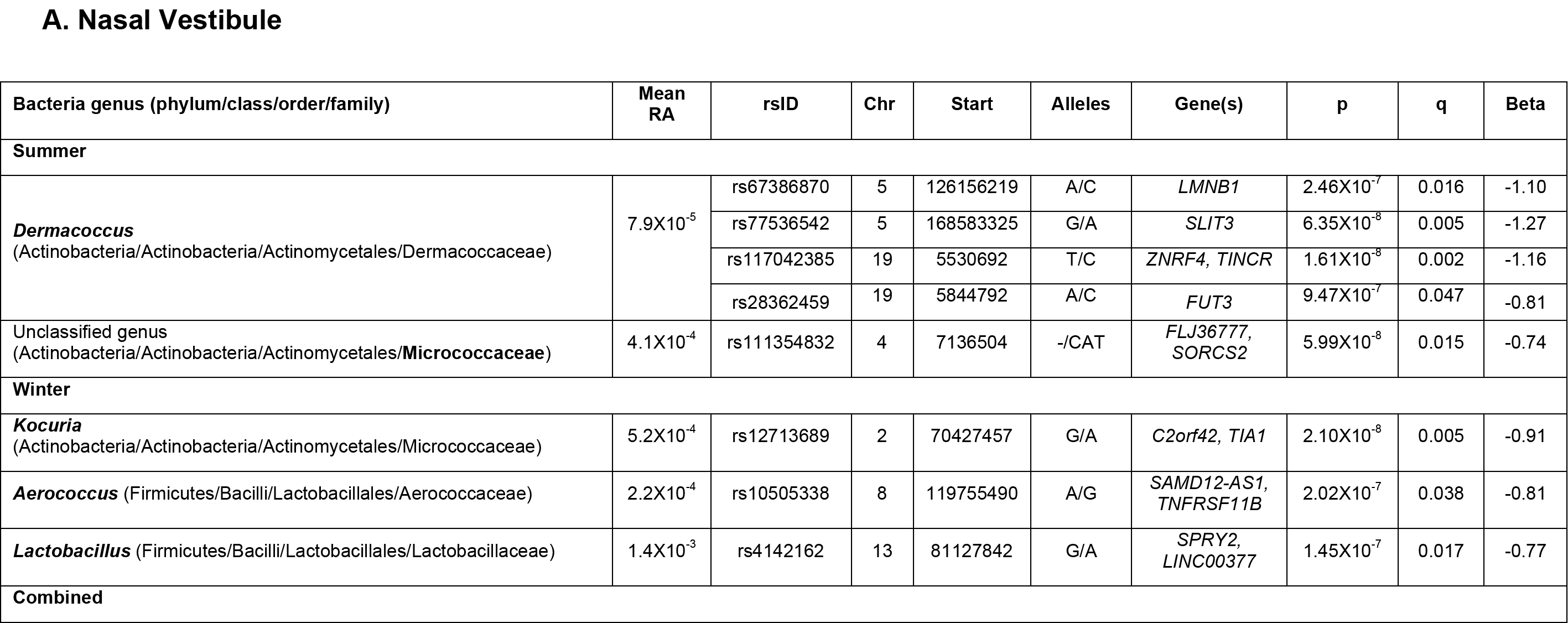
QTL mapping results of nasal microbiome relative abundance. **A. Nasal vestibule.** 14 host variants were associated at a q < 0.05 with the relative abundance of 10 genera. rs111354832 is associated with an unclassified genus of family Micrococcaceae in the summer and in the combined sample. **B. Nasopharynx.** 23 host variants were associated at a q < 0.05 with the relative abundance of 14 genera. At this site 2 SNPs (rs1653301 and rs7702475) are associated with more than one bacterium. rsIDs presented for dbSNP142. Alleles presented as minor/major. Direction of effect is presented for the minor allele. When genus level is unclassified, highest classified taxonomic level is bolded. RA, relative abundance; Chr, chromosome.

|  |  |  |  |  |  |  |  |  |  |
| --- | --- | --- | --- | --- | --- | --- | --- | --- | --- |
| Unclassified genus<br>(Actinobacteria/Actinobacteria/Actinomycetales/**Intrasporangiaceae**) | 9.2X10<sup>-5</sup> | rs11085969 | 19 | 15792546 | A/G | *CYP4F12* | 5.84X10<sup>-8</sup> | 0.005 | -0.91 |
| **Microbispora**<br>(Actinobacteria/Actinobacteria/Actinomycetales/Micrococcaceae) | 5.5X10<sup>-4</sup> | rs2891405 | 12 | 113152097 | G/A | *MIR1302-1,*<br>*RPH3A* | 1.50X10<sup>-7</sup> | 0.038 | -0.81 |
| Unclassified genus<br>(Actinobacteria/Actinobacteria/Actinomycetales/**Micrococcaceae**) | 3.5X10<sup>-4</sup> | rs111354832 | 4 | 7136504 | -/CAT | *FLJ36777,*<br>*SORCS2* | 3.45X10<sup>-8</sup> | 0.017 | -0.77 |
| **Peptoniphilus** (Firmicutes/Clostridia/Clostridiales/Tissierellaceae) | 8.9X10<sup>-5</sup> | rs9865782 | 3 | 113652774 | A/G | *GRAMD1C* | 1.17X10<sup>-7</sup> | 0.005 | -0.91 |
| **Paracoccus**<br>(Proteobacteria/Alphaproteobacteria/Rhodobacterales/Rhodobacteraceae) | 6.5X10<sup>-5</sup> | rs9953410 | 18 | 29532946 | C/A | *TRAPPC8,*<br>*RNF125* | 2.21X10<sup>-7</sup> | 0.038 | -0.81 |
| **Enterobacter**<br>(Proteobacteria/Gammaproteobacteria/Enterobacteriales/Enterobacteriaceae) | 6.4X10<sup>-5</sup> | rs11042877 | 11 | 10576232 | A/C | *MRVI1-AS1* | 3.86X10<sup>-7</sup> | 0.017 | -0.77 |
|  |  | rs12446497 | 16 | 7341674 | A/G | *RBFOX1* | 5.84X10<sup>-7</sup> | 0.005 | -0.91 |

B. Nasopharynx
| Bacteria species (phylum/class/order/family) | Mean RA | rsID | Chr | Start | Alleles | Gene(s) | p | q | Beta |
| --- | --- | --- | --- | --- | --- | --- | --- | --- | --- |
| Summer |  |  |  |  |  |  |  |  |  |
| ***Aerococcus*** (Firmicutes/Bacilli/Lactobacillales/Aerococcaceae) | 6.5X10<sup>-4</sup> | rs7702475 | 5 | 58088523 | A/G | *RAB3C* | 3.97X10<sup>-8</sup> | 0.010 | 0.77 |
| Unclassified genus (Firmicutes/Bacilli/Lactobacillales/**Aerococcaceae**) | 5.3X10<sup>-4</sup> | rs11888528 | 2 | 120118764 | C/T | *C2orf76* | 1.43X10<sup>-7</sup> | 0.038 | 0.79 |
| ***Methylobacterium***<br>(Proteobacteria/Alphaproteobacteria/Rhizobiales/Methylobacteriaceae) | 4.3X10<sup>-4</sup> | rs308961 | 3 | 12150014 | T/G | *SYN2* | 8.12X10<sup>-7</sup> | 0.034 | -0.70 |
|  |  | rs10547084 | 4 | 37753111 | -/TCTC | *RELL1,PGM2* | 4.93X10<sup>-7</sup> | 0.031 | 0.77 |
|  |  | rs67737950 | 4 | 40260058 | G/C | *RHOH, LOC101060498* | 2.73X10<sup>-7</sup> | 0.023 | 0.85 |
|  |  | rs7702475 | 5 | 58088523 | A/G | *RAB3C* | 2.46X10<sup>-7</sup> | 0.023 | 0.77 |
|  |  | rs1278260 | 10 | 127731197 | C/A | *ADAM12* | 6.21X10<sup>-7</sup> | 0.031 | 0.95 |
| Unclassified genus<br>(Proteobacteria/Alphaproteobacteria/Caulobacterales/**Caulobacteraceae**) | 8.5X10<sup>-4</sup> | rs927984 | 6 | 25412987 | T/C | *LRRC16A* | 2.78X10<sup>-7</sup> | 0.036 | 0.94 |
|  |  | rs1543603 | 6 | 25413922 | A/G | *LRRC16A* | 2.30X10<sup>-8</sup> | 0.006 | 0.88 |
| Winter |  |  |  |  |  |  |  |  |  |
| ***Mycobacterium***<br>(Actinobacteria/Actinobacteria/Actinomycetales/Mycobacteriaceae) | 2.2X10<sup>-4</sup> | rs1802665 | 10 | 61788623 | G/T | *ANK3* | 8.73X10<sup>-8</sup> | 0.007 | -0.98 |
| ***Gemella*** (Firmicutes/Bacilli/Gemellales/Gemellaceae) | 3.7X10<sup>-4</sup> | rs17631306 | 1 | 111072322 | A/G | *KCNA10, KCNA2* | 2.94X10<sup>-8</sup> | 0.008 | 1.27 |
| Unclassified genus (Firmicutes/Bacilli/**Lactobacillales**) | 5.7X10<sup>-4</sup> | rs1153741 | 2 | 182860422 | G/A | *PPP1R1C* | 1.52X10<sup>-7</sup> | 0.039 | -0.83 |
| **Combined** |  |  |  |  |  |  |  |  |  |
| **Gordonia**<br>(Actinobacteria/Actinobacteria/Actinomycetales/Gordoniaceae) | 1.1X10<sup>-4</sup> | rs61925863 | 12 | 66694722 | C/G | *IRAK3, HELB* | 2.88X10<sup>-7</sup> | 0.038 | -0.90 |
|  |  | rs12435212 | 14 | 85483485 | G/T | *NONE, LINC00911* | 1.93X10<sup>-8</sup> | 0.005 | 0.88 |
| Unclassified genus (Firmicutes/Clostridia/Clostridiales/**Clostridiaceae**) | 1.0X10<sup>-2</sup> | rs9661504 | 1 | 205915667 | A/T | *SLC26A9, FAM72C* | 4.16X10<sup>-7</sup> | 0.046 | -0.70 |
|  |  | rs10232599 | 7 | 46035291 | G/A | *IGFBP3, TNS3* | 1.14X10<sup>-7</sup> | 0.029 | -0.67 |
| Unclassified genus (Actinobacteria/Actinobacteria/**Actinomycetales**) | 1.3X10<sup>-3</sup> | rs12156316 | 8 | 41706484 | T/C | *ANK1* | 8.16X10<sup>-7</sup> | 0.042 | 0.58 |
|  |  | rs10901086 | 9 | 134635034 | T/C | *RAPGEF1, MED27* | 7.05X10<sup>-7</sup> | 0.042 | 0.89 |
|  |  | rs12244238 | 10 | 6083239 | G/A | *IL2RA* | 6.82X10<sup>-7</sup> | 0.042 | -0.69 |
| Unclassified genus<br>(Actinobacteria/Actinobacteria/Actinomycetales/**Micrococcaceae**) | 5.0X10<sup>-4</sup> | rs3006458 | 1 | 153320372 | T/G | *PGLYRP4* | 5.10X10<sup>-7</sup> | 0.032 | -0.75 |
|  |  | rs4774283 | 15 | 58114121 | T/G | *GCOM1, ALDH1A2* | 2.56X10<sup>-7</sup> | 0.032 | -0.57 |
|  |  | rs4814474 | 20 | 16322199 | A/C | *KIF16B* | 9.35X10<sup>-7</sup> | 0.040 | 0.56 |
| **Rhodococcus**<br>(Actinobacteria/Actinobacteria/Actinomycetales/Nocardiaceae) | 6.4X10<sup>-5</sup> | rs1653301 | 2 | 201076401 | A/G | *C2orf47, SPATS2L* | 1.45X10<sup>-8</sup> | 0.004 | -0.82 |
| Unclassified genus<br>(Actinobacteria/Actinobacteria/Actinomycetales/**Sporichthyaceae**) | 1.5X10<sup>-4</sup> | rs13128830 | 4 | 21455808 | T/C | *KCNIP4* | 3.03X10<sup>-7</sup> | 0.022 | 0.64 |
| Unclassified genus<br>(Proteobacteria/Alphaproteobacteria/Sphingomonadales/**Sphingomonadaceae**) | 5.1X10<sup>-3</sup> | rs1653301 | 2 | 201076401 | A/G | *C2orf47, SPATS2L* | 9.48X10<sup>-8</sup> | 0.024 | -0.80 |

The most significant association was with an intergenic SNP 8kb upstream of the *TINCR* gene on chromosome 19 and the abundance of *Dermacoccus* (phylum Actinobacteria) in the nasal vestibule in the summer (rs117042385; p = 1.61⨯10^−8^; q = 0.002, Figure 4). *TINCR* is a long non-coding RNA gene that controls human epidermal differentiation and directly binds to the peptidoglycan recognition protein 3 (*PGLYRP3*) transcript (Kretz et al. 2013). A second mbQTL (rs28362459), located 314 kb downstream from the *TINCR* mbQTL (*r*^2^ = 0.26; D’=0.76), was also associated with the RA of *Dermacoccus* in the same site and season as *TINCR* (p = 9.47⨯10^−7^, q = 0.047, Figure 4). rs28362459 is a missense variant in fucosyltransferase 3 (*FUT3*), a gene essential for the synthesis of Lewis blood groups (Taylor-Cousar et al. 2009; Yamamoto et al. 2014).

To determine if the association of increased *Dermacoccus* RA with the rs28362459-C allele in *FUT3* is independent of the association with the rs117042385-C allele upstream of *TINCR*, we phased the two variants and examined the four haplotypes (seven diplotypes) present in our sample. This revealed independent effects of genotypes at both SNPs contributing to the RA of *Dermacoccus* (p = 4.65 ⨯10^−9^; Figure 4C). In particular, individuals who were homozygous for both alleles (rs117042385-CC/rs28362459-CC) had the highest RA of *Dermacoccus*, while one or two copies of the *FUT3* rs28362459-A allele on a homozygous *TINCR* rs117042385-CC background was associated with decreased RA. Overall, the presence of a *FUT3* rs28362459-A allele was associated with lower RA regardless of genotype at *TINCR* rs117042385. The two individuals who were homozygous for both the *TINCR* rs117042385-T and *FUT3* rs28362459-A alleles did not have any *Dermacoccus* sequences detected. Overall, these results suggest that the *FUT3* rs28362459 and *TINCR* rs117042385 variants (or variants in strong LD with them) are exerting independent effects on the RA of *Dermacoccus*.

**Figure 4:**
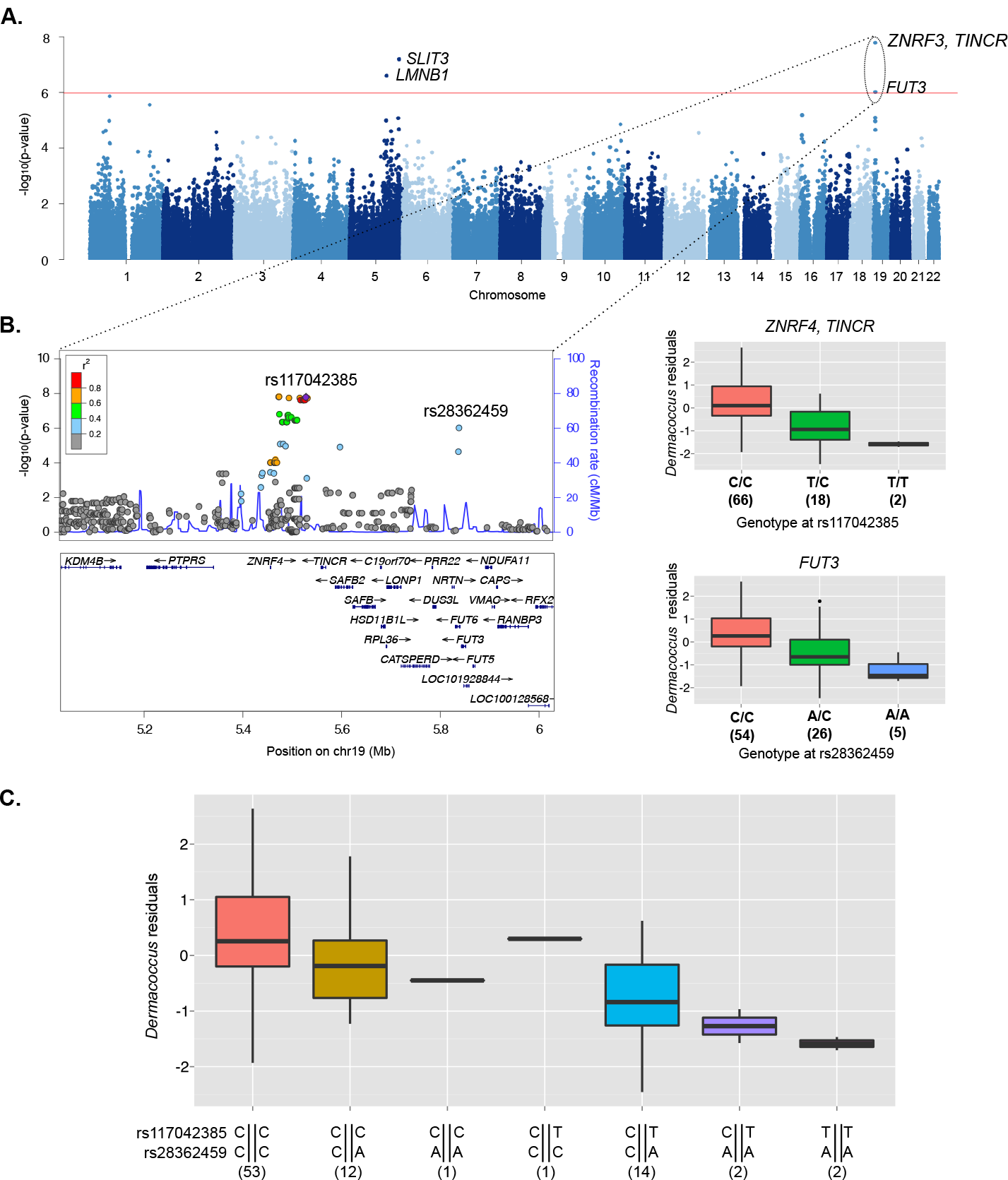
Associations with the RA of *Dermacoccus* in the nasal vestibule in the summer. **A. Manhattan plot.** Association results are presented for variants pruned for LD (r^2^ < 0.5). Four variants on chromosomes 5 and 19 are associated with the RA of *Dermacoccus* at a q < 0.05 significance threshold (red line). **B. Locus and genotype plots for the 2 mbQTLs on chromosome 19.** Variants included in the locus plot are those with MAF > 10% in the Hutterites, prior to LD pruning. Genotype plots show both minor alleles (T at rs117042385 and A at rs28362459) are associated with lower *Dermacoccus* RA. **C. Boxplots of *Dermacoccus* residuals for rs117042385 and rs28362459 phased haplotypes.** Numbers underneath each boxplot represent the number of individuals in each genotype or haplotype class.

Another mbQTL that linked PGLYRP genes more directly to host regulation of the microbiome is an association between a missense variant in *PGLYRP4* (rs3006458) on chromosome 1 and the RA of an unclassified genus of family Micrococcaceae (phylum Actinobacteria) in the combined season nasopharynx sample (p = 5.10⨯10^−7^; q = 0.032, Supplemental Figure S4A). This same SNP was also associated with genus *Aerococcus* in the nasopharynx in the winter at a less stringent q value cutoff (phylum Firmicutes; p = 1.28⨯10^−6^; q = 0.06). Peptidoglycan recognition proteins (PGRPs) are a conserved family of antibacterial pattern recognition molecules that directly bind peptidoglycan and other bacterial cell wall components, including lipopolysaccharide (LPS) (Kashyap et al. 2011).

### *mbQTL* associations *with multiple bacteria*

Five mbQTLs (q < 0.05), including the *PGLYRP4* mbQTL discussed above, had associations with multiple bacteria within the same nasal site at a relaxed significance threshold (q < 0.10). The largest number of associations identified with a single mbQTL was an intronic variant in the Leucine Rich Repeat Containing 16A (*LRRC16A;* rs1543603) and the RAs of five Proteobacteria in the nasopharynx in the summer (unclassified genus of family Caulobacteracea, unclassified genus of family Bradyrhizoviaceae, *Parvibaculum, Blastomonas* and *Rheinheimera)*, of which Caulobacteraceae was the most significant (p = 2.30x10^−8^, q = 0.006; Supplemental Figure S4B). *LRRC16A* encodes CARMIL (capping protein, Arp2/3, and Myosin-I linker), a protein that plays an important role in cell shape and motility (Yang et al. 2005).

The association between genotype at a single SNP, rs1543603, with the RAs of five genus level bacteria suggested potential functional community level relationships between these five Proteobacteria. Indeed, the RAs of all five Proteobacteria were correlated with each other (correlation coefficients > 0.773; median 0.924). A co-occurrence network (Friedman and Alm 2012) assigned all five bacteria to a single network that included 13 bacteria (12 Proteobacteria and one Bacteroidetes) from among the 90 genera tested in the nasopharynx in the summer (Figure 5). In this network, genera from families Bradyrhizobiaceae and Caulobacteraceae, two of the five bacteria associated with rs1543603 (p = 3.25⨯10^−7^ and 2.30⨯10^−8^, respectively) are the largest hubs with nine neighbors each. These findings suggest that host genetic effects can act to modulate microbial community patterns, by directly affecting host-microbe interactions with only one or a few main drivers of the community.

**Figure 5:**
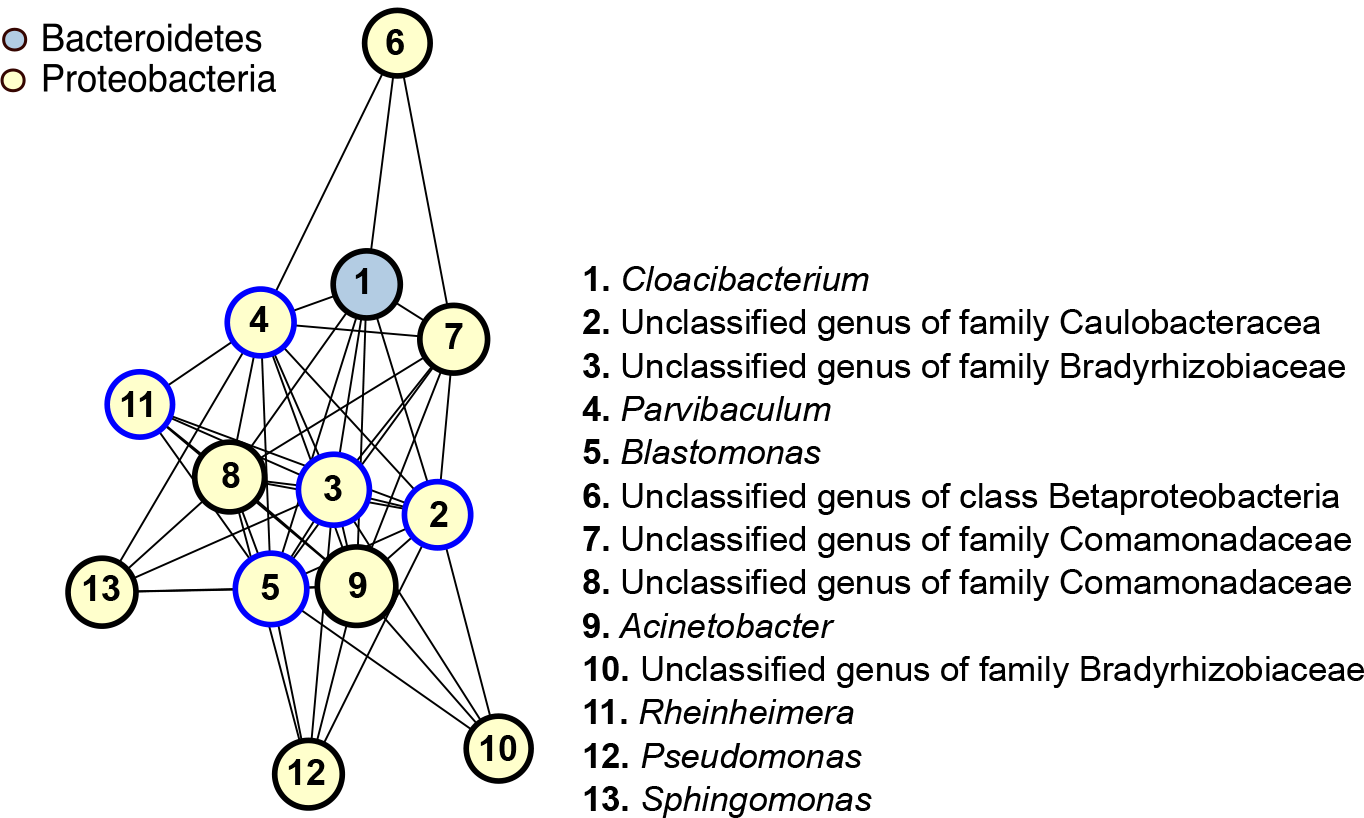
Five genus-level bacteria associated with rs1543603 are hubs in a cooccurrence module of 12 Proteobacteria and one Bacteroidetes. Co-occurrence networks built from correlation coefficients between all 90 genus level RAs determined in the nasopharynx summer sample. Nodes represent bacteria and are listed by number, colored by phylum and sized proportionally to the RA of each bacterium. Edges represent correlations greater than 0.75. Blue node borders represent the five bacteria associated with rs1543603, an intronic variant in *LRRC16A*.

### Pathway analyses of genes near mbQTLs

To further understand how host genetic variation regulates nasal microbiome composition and to identify shared pathways among the mbQTLs identified in this study, we selected the closest gene to each mbQTL (q < 0.10; 131 genes) and to all variants in LD (*r*^2^ > 0.8) with each mbQTL, using LD estimates in the Hutterites. We then generated protein-protein interaction networks among these genes, using Ingenuity Pathway Analysis Knowledge Base (IPA^®^, QIAGEN Redwood City, CA), a curated database of biological interactions and functional annotations. IPA identified two networks with Fisher exact p < 10^−25^ (Figure 6). The most significant network included 21 of the 131 genes, nine of which were near mbQTLs with q < 0.05 (Fisher exact p = 10^−43^; Figure 6A). This network contained many hubs including *SMAD2*, a gene that regulates the production of immunoglobulin A (IgA) by LPS-activated B-cells and activates immune response at other mucosal surfaces upon stimulation by pathogenic microbes (Malhotra and Kang 2013). The second significant network (Fisher exact p = 10^−29^) contained 17 of the 131 genes, also with nine genes near mbQTLs with q < 0.05. Many of the hubs in this network represent important modulators of mucosal immunity, including immunoglobulins A and G (IgG and IgG2a), IL12/IL12RA, TCR and STAT5A/B (Macpherson et al. 2008; Holt et al. 2008; Mantis and Forbes 2010).

**Figure 6:**
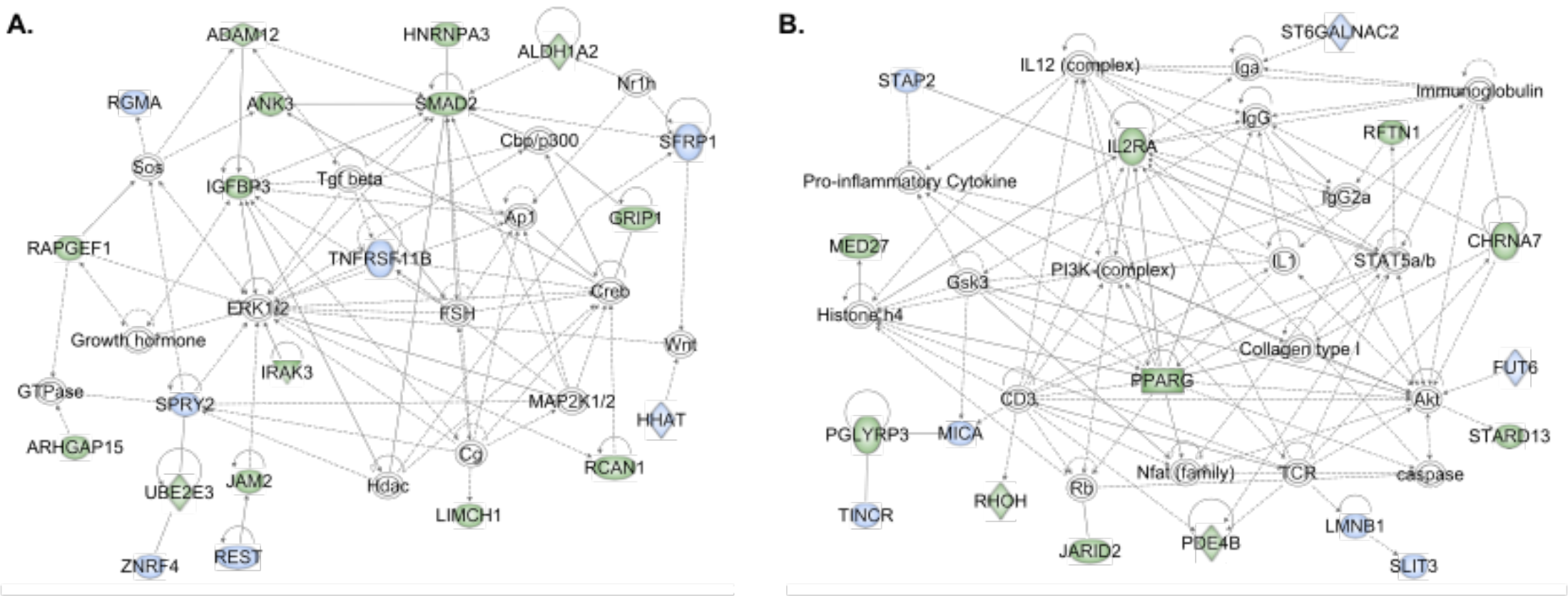
Ingenuity Pathway Analysis (IPA) interaction networks. Networks show genes near nasal mbQTLs are enriched for mucosal immunity pathways. Two significant networks (p < 10^−25^) are presented. **A.** Network one is centered on *SMAD2* and *ERK1/2* (p < 10^−43^; 21 genes).**B.** Network two is a highly connected network centered on *IL2RA*, STAT5a/b and *IL12*, among others. This network contains many of the key regulators of mucosal immunity (p < 10^−29^; 17 genes). Node color represents genes near microbiome QTL associations in the nasal vestibule (blue) or in the nasopharynx (green); open symbols are genes added by IPA. Edges represent direct (solid) and indirect (dashed) interactions in the IPA Knowledge Base database. Node shapes correspond to functional classes of gene products: concentric circles for groups or complexes, diamonds for enzymes, rectangles for transcriptional regulators or modulators, ovals for trans-membrane receptors and circles for other.

### Comparison of genes near mbQTLs in the Hutterites to the TwinsUK study

The TwinsUK study, the largest human gut microbiome QTL study to date, recently reported a candidate gene study (Goodrich et al. 2016) in which 17 genes within 10 kb of 15 mbQTLs were associated with 15 different taxa (false discovery rate (FDR) less than 5%). To determine if genes associated with the RA of bacteria in the upper airway also influence the RA of bacteria in the gut, we compared 53 genes near the 37 mbQTLs discovered in our study (q < 0.05) to the 17 genes reported in the TwinsUK Study. Two different intronic variants in the slit guidance ligand 3 (*SLIT3*) gene on chromosome 5 were associated with the RA of an unclassified genus of family *Clostridiaceae* in TwinsUK (rs10055309) and with the RA genus *Dermacoccus* in the nasal vestibule in the summer in our study (rs77536542; p = 6.35⨯10^−8^; Figure 4A). SNP rs10055309 was the most significant QTL reported the TwinsUK study. *SLIT3* is a secreted protein that is widely expressed across many tissues with highest expression in skin, brain cerebellum and lung (Dickinson et al. 2004). *SLIT3* hypermethylation has been reported in a number of human cancers (Dickinson et al. 2004) and *SLIT3* expression is increased in LPS stimulated macrophages in mice (Tanno et al. 2007). The combined data from the TwinsUK Study and our study suggests that this gene may play a role in the modulating bacterial abundances across diverse body sites in humans.

## DISCUSSION

Our study is the first to assess the role of genome-wide host genetic variation in shaping the human microbiome at two upper airway sites. We first demonstrated reduced bacterial beta diversity between more closely related pairs of individuals, and then discovered associated genetic variants at functionally related genes. These combined results indicate a significant role for host genotype in patterning microbial diversity in the nose.

Our results further suggest that the upper airway may be the site of important gene-environment interactions. In this context, host genotype at many loci may ultimately impact health and disease by modulating particular members of the microbial community. For example, a missense variant in fucosyltransferase 3 (*FUT3*; rs28362459) was strongly associated with decreased abundance of *Dermacoccus* in the nasal vestibule in the summer. This SNP is predicted to be deleterious by both Polymorphism Phenotyping (PolyPhen) v2 (Adzhubei et al. 2010) and Combined Annotation Dependent Depletion (CADD) (Kircher et al. 2014) scores (0.997 and 15.12, respectively). Interestingly, the non-secretor phenotype, characterized by a null variant in another FUT gene, *FUT2*, and the resulting absence of ABH antigens in the mucosa in homozygotes for the null allele, influences the composition and diversity of the microbiome in the human intestinal tract (Wacklin et al. 2011; Rausch et al. 2011). Moreover, variants in both *FUT2* and *FUT3* have been shown in GWAS to increase susceptibility to diseases associated with both mucosal surface pathobiology and microbiome composition, such as cystic fibrosis (Taylor-Cousar et al. 2009), Crohn’s disease (Hu et al. 2014) and ulcerative colitis (Hu et al. 2016). Our study extends a role for fucosyltransferases to the nasal mucosal surface, and further implicates host genetic influences on bacterial diversity at this site.

Four mbQTLs show effects on phylogenetically diverse phyla and two were identified in different seasons and nasal sites. In particular, a missense variant in *PGLYRP4* (rs3006458) was associated with the abundance of genus *Aerococcus* (Firmicutes) in the nasopharynx in the summer and with family Micrococcacea (Actinobacteria) in the nasopharynx in the combined sample. The RAs of *Aerococcus* and Micrococcacea are only weakly correlated with each other indicating that these are likely independent associations. Moreover, the associations with phylogenetically distant bacteria and in different subsamples (summer vs. combined) suggest that *PGLYRP4* has pleiotropic effects over several organisms. Alternatively, the SNP identified in our study (rs3006458) could be tagging a haplotype with multiple variants that have independent effects on different bacterial abundances. The genomic region that includes the PGLYP genes includes a cluster of genes implicated in epidermal barrier function (Toulza et al. 2007) and SNPs in this region show extensive LD. However, independent evidence suggests that *PGLYRP4* may be the target gene of this association. The rs3006458-T allele, which is associated with lower RA of *Aerococcus* and Micrococcacea in our study, was associated with increased gene expression of the *PGLYRP4* gene in epithelial and mucosal tissues (skin, small intestine and esophageal mucosa) in the Genotype-Tissue Expression (GTEx Consortium 2015), and in lung tissue in a separate eQTL study (Hao et al. 2012). These tissues serve as physical barriers and provide innate immune functions essential for antimicrobial defense (Gallo and Hooper 2012). Collectively, these data suggest that host genotype at rs3006458 (or a variant in LD with rs3006458) regulates the expression in *PGLYRP4* in the skin, lung and airway mucosa and functions to modulate bacterial abundance, possibly beyond the two genera identified in this study. The link between genetic variation in host PGRPs and microbiome abundance revealed in this study indicates that at least some of the important role these proteins play in modulating communities of symbiotic organisms (Royet et al. 2011) is attributable to host genetic variation.

Eight of the mbQTLs identified in our study at a relaxed q < 0.10 influenced the abundance of more than one organism. Most were identified within the same season and nasal site and four influenced the abundance of multiple closely related bacteria. For example, an intronic variant in *LRRC16A* (rs1543603) was associated with the abundance of five highly correlated genera of phylum Proteobacteria in the nasopharynx in the summer. These five bacteria co-occur in a larger network of 12 Proteobacteria and one Bacteroidetes, suggesting that they may be physically interacting and that their overall community structure is influenced by host genotype. Although not much is known about *LRRC16A*, other proteins with leucine-rich repeat (LRR) domains, such as nucleotide-binding oligomerization domain receptors (NODs) and toll-like receptors (TLRs) (Ng et al. 2011), function as recognition receptors in innate immunity.

Although our study provides novel insights into host genetic influences on the nasal microbiome, there are some limitations. In particular, the size of our sample is relatively small for genetic mapping (77–88 individuals in seasonal analyses; 125 and 133 individuals in the combined). We reasoned that the reduced environmental heterogeneity among Hutterite individuals would enhance the effects of genetic variation and facilitate the detection of associated variants. While we were successful in identifying mbQTLs, we acknowledge that there are likely many more associations to be found in larger samples. A second limitation is the multiple testing burden that results from the high dimensionality of the microbiome. While we reduced the number of tests performed by mapping only genus level bacteria present in the majority of individuals, we only corrected for multiple testing within each study and did not correct for the 90 bacteria for which we performed mbQTL mapping. Although we used a fairly stringent threshold of genome-wide significance (q < 0.05), we acknowledge that some of our findings may be false positives. Lastly, the genetic effects revealed by our study are context specific due to the many environmental and stochastic factors that affect microbiome composition and, therefore, challenging to replicate. For example, even within our study of a relatively homogenous population, we detected significant effects of season, age, and gender. In fact, most of the mbQTLs that we identified were specific to one season and demonstrate that even small temporal changes (~6 months) in the RAs of bacteria within the same individuals can mask or enhance genetic effects. Although we did not formally replicate the results in independent populations, the identification of intronic variants within the gene *SLIT3* in our study and in the TwinsUK study (Goodrich et al. 2016) bolsters confidence in the involvement of this gene in regulating microbial structure across multiple mucosal sites.

These limitations aside, our study provides evidence for genetic contributions to modulating variability of the nasal microbiome, a trait that has been linked to a number of airway diseases (Dickson et al. 2014). Importantly, our findings support the concept that host genetic variation directly influences the expression or function of genes that are specifically involved in innate mucosal immunity pathways. Such a framework is consistent with previous reports showing that antimicrobial peptides (Salzman et al. 2010; Royet et al. 2011) and host immunity (Hooper and Macpherson 2010) are key modulators of microbial defense in the mucosa. Our data further suggest that host genetic effects on immune genes modulate particular bacteria or the structure of whole microbial communities in the upper airways. We speculate that interactions between host genetics and microbiome structure or composition in the upper airway can influence dysbiotic tendencies that may predispose to respiratory disease and could be subject to intervention. Indeed, moving forward, more detailed analyses of the complex relationship between genetic variation in host mucosal immunity and the microbiome — captured here in a snapshot in the upper airway — are required to fully characterize determinants of an inherently dynamic microbial ecosystem. Such work could potentially identify targets for novel therapeutic strategies useful across a wide range of respiratory diseases.

## METHODS

### Sample collection

Nasal brushings from Hutterites ages 16 to 78 from five colonies located in South Dakota, all within 14 miles of each other, were collected at two time points, winter (January/February 2011) and summer (July 2011), and from two nasal sites, the nasal vestibule and the nasopharynx. Samples from each of the two nasal sites were collected from opposite nares using sterile flock collection swabs (Puritan^©^ 25-3316). The Human Microbiome Project anterior nare collection protocol (Aagaard et al. 2013) was used for the nasal vestibule and an adaptation of the Pasculle et al. protocol (Pasculle et al. 2008) was implemented for the nasopharynx. After excluding samples with low DNA yield, low sequencing read depth, antibiotic history within the prior 3 months, or missing genotypes, our final data consists of 133 individuals with nasal vestibule samples (87 summer and 80 winter) and 125 individuals with the nasopharynx samples (88 summer and 77 winter; Table 1).

### Sample DNA extraction library preparation and sequencing

Nasal brushes were immediately frozen at −20°C following collection, shipped on dry ice, and stored at −80°C. DNA extraction was carried out using the BiOstic^®^ Bacteremia DNA Isolation Kit (12240-50). DNA concentration and purity were assessed using the Nanodrop 1000 spectrophotometer (Thermo Scientific, IL, USA). The 16S rRNA gene V4 region was amplified following conditions in Caporaso et al. protocol (Caporaso et al. 2012) using 62 different region-specific primers labeled with a unique 12-base Golay barcode sequence in the reverse primer. Final libraries were quality controlled prior to pooling with the Agilent Bioanalyzer DNA 1000 (Agilent Technologies, CA, USA). Libraries were pooled into 8 pools of 62 samples and sequenced on the HiSeq2000 platform (Illumina Inc, CA, USA) under a single end 102 base pair protocol.

### Sequencing and taxonomic classification

Data was pre-processed using CASAVA 1.8.1. Following sample de-multiplexing, 617,909,462 sequence reads were processed using the Quantitative Insights into Microbial Ecology (QIIME)
1.8.0 toolkit (Caporaso et al. 2010b). Quality controlled reads were required to have an exact match to an expected barcode, zero ambiguous base calls, less than three consecutive low quality base calls, and a minimum Phred quality score of 20 along the entire read. We used an open-reference OTU workflow where sequences were first clustered against the Greengenes May 2013 reference (Caporaso et al. 2010b; DeSantis et al. 2006) and reads that did not cluster with known taxa (97% identity) were subjected to *de novo* clustering. Representative sequences were aligned using PyNAST version 1.2.2 (Caporaso et al. 2010a) and the taxonomy of each OTU cluster was assigned with the *uclust* classifier version 1.2.22q (Edgar 2010). We applied an OTU abundance filter of 0.005% (Bokulich et al. 2013) to reduce spurious OTUs and obtained a final dataset of 563 OTUs.

### Data processing

#### *Seasonal*

For each of the four seasonal groups (nasal vestibule in the summer and in the winter, and nasopharynx in the summer and winter), a genus level RA table was calculated after subsampling reads to 250,000 per sample. Each bacteria’s RA was then quantile normalized using the *qqnorm* function in R. Next, PCA was performed using the *prcomp* function in R and each of the top 10 principal components (PCs, explaining ~76%–78% of the variance) were tested in a linear model against technical covariates. In at least one of the seasonal groups, we identified correlations between one of the PCs with DNA concentration prior to PCR, final base pair fragment size and date of sampling (p < 0.001). PCR adapter barcode, library batch, order within library batch, and final library concentration were not significant. After regressing out the identified technical covariates from the normalized RAs, we performed PCA on the residuals and tested for associations between the top 10 PCs and biological covariates. Age and sex were significant and were therefore adjusted for in all subsequent analyses. Next, to reduce the burden of multiple testing in the seasonal mapping studies, we removed genera that were detected in fewer than 75% of individuals. This resulted in 78 genus level RAs in the nasal vestibule in the summer, 52 in the nasal vestibule in the winter, 90 in the nasopharynx in the summer and 59 in the nasopharynx in the winter.

#### *Combined seasons*

Although combining samples across seasons could introduce noise, it provides the largest possible sample size and consequently greatest power for genetic associations with bacteria that do not vary in abundance across seasons. Therefore, for each nasal site, we averaged the summer and winter genus level RA residuals obtained after quantile normalization and the regression of identified technical covariates for individuals with measurements during both seasons, or included the one season result for those with only one measurement (referred to as the combined sample). We selected all genus level bacteria present in at least 75% of individuals in either season, which resulted in 76 genus level RAs in the nasal vestibule and 90 in the nasopharynx. We performed PCA on the combined seasons matrix to verify variation among samples did not separate the combined samples from the samples with one seasonal measurement. Season of origin (summer, winter or averaged) was not correlated with any of the top 10 principle components in either nasal site (Supplemental Figure S5).

### Genotype data

The Hutterite individuals in our study are related to each other in a 13-generation pedigree that includes 3,671 individuals, all of whom originate from 64 founders. Using PRIMAL (Livne et al. 2015), an in-house pedigree-based imputation algorithm, whole genome sequences from 98 Hutterite individuals were phased and imputed to 1,317 Hutterites who were previously genotyped on Affymetrix arrays (Ober et al. 2008; Yao et al. 2014; Cusanovich et al. 2012). For mapping studies, we first selected 3,161,460 variants with genotype call rates greater than 95% in our sample and minor allele frequencies (MAF) > 0.10 in any of the 4 season/site subsamples. Next, we estimated LD in the Hutterite data using PLINK (Purcell et al. 2007), and pruned variants for LD using an r^2^ threshold of 0.5, to yield a final set of 148,653 variants for mapping studies.

### Diversity metrics

Alpha diversity metrics at the species level (observed species, Shannon index and evenness) were calculated in QIIME (Caporaso et al. 2010b) using the alpha_diversity.py script after subsampling reads from 1,000 to 10,000 every 1,000 reads, from 10,000 to 100,000 every
10.0 reads and from 150,000 to 250,000 every 50,000 reads. Each subsampling series was completed 10 times and rarefaction curves were plotted. Diversity metrics were averaged from the 250,000 read subsamples using the collate_alpha.py script and this metric was compared across seasons and nasal sites.

To calculate beta diversity, we first obtained the OTU table using *phyloseq* (McMurdie and Holmes 2013), quantile normalized OTU abundances using *qqnorm* in R and regressed out technical covariates (DNA concentration prior to PCR, final base pair fragment size and date of sampling). Next, we calculated pairwise Euclidean distance using the *vegdist* function in the *vegan* R package.

### Kinship associations to beta diversity

Pair-wise kinship coefficients were previously calculated by PRIMAL (Livne et al. 2015) using 271,486 variants genotyped on Affymetrix platforms. The average kinship coefficient between all pairs of individuals (n = 144) in our study was 4.51% (range 0.60%–32.03%). We performed 10.0 permutations to asses the association between pairwise Euclidian distances and kinship coefficients in the combined seasons samples (nasal vestibule 8,778 pairs, nasopharynx 7,750). The p-value is the number of times out of 10,000 permutations that the Spearman correlation of the permuted sequence pair was more extreme than the observed pair.

### Co-occurrence network analyses

We used SparCC (Friedman and Alm 2012) to calculate nasopharynx in the summer correlation coefficients between all 90 genera tested in our mapping studies. We applied default settings and assigned p-values calculated from 100 bootstraps. Co-occurrence networks were generated from the SparCC correlation matrix for genera with correlation *r^2^* > 0.75 and p < 0.01 (1/100 bootstraps). The network was generated using *igraph* R package, where nodes represent each genera and edges represent correlations between the genera above the applied threshold.

### Ingenuity Pathway Analysis (IPA) of protein-protein interaction networks

We selected the closest gene to 1,413 variants (131 genes) with Hutterite linkage disequilibrium (LD) *r^2^* > 0.8 with the 108 mbQTLs (q < 0.10). To interrogate and visualize network associations, we used the Ingenuity Pathway Analysis Knowledge Base (IPA^®^, QIAGEN Redwood City, CA), limiting interactions to primary cells or tissues. The network scores generated by IPA are based on a right-tailed Fisher’s Exact test comparing the observed and expected mbQTL genes present in a pathway relative to the IPA database.

## DATA ACCESS

Submission to dbGaP is currently in progress.

## ACKNOWLEDGEMENTS

We would like to thank members of the Hutterite community for their continuous participation in our studies. We would also acknowledge members of the Ober, Gilbert and Gilad labs for useful discussions; Sally Cain and Rebecca Anderson for assistance with sample collection; Amy Mitrano for library preparation; James Lane for technical support; and Jack Gilbert for analysis advice.

This work was supported by NHLBI (HL085197), the NIAID (AI106683 and AI095230), NIA (AG036762) and the AAAAI (Allergy, Asthma, and Immunology Education, and Research Trust award). C.I. was supported by National Institutes of Health Grant T32 GM007197 and by the Ruth L. Kirschstein National Research Service Award (HL123289).

## AUTHOR CONTRIBUTIONS

CO, JP and ED collected samples. ED generated sequencing libraries. CI, YG, DN, CO and JP designed the study and performed analyses. CI, CO and JP and wrote the manuscript. All authors discussed results and approved the manuscript.

